# “Biomarkers of recovery: characterizing trophic flow following ecological restoration”

**DOI:** 10.1101/2024.11.14.623625

**Authors:** Nathan B. Spindel, Aaron W. E. Galloway, Julie B. Schram, Gwiisihlgaa Daniel Mcneill, Sgiids Kung Vanessa Bellis, Niisii Guujaaw, Jaasaljuus Yakgujanaas, Markus Thompson, Lynn C. Lee, Daniel K. Okamoto

## Abstract

Coastal kelp forests are important sources of primary productivity and provide essential habitat and ecosystem services. In many areas around the world, the formation and persistence of urchin barrens threatens kelp forest ecosystems. Over the past several decades, restoration efforts have emerged aiming to increase the abundance of foundation species like kelp in such systems. However, we lack a comprehensive understanding of how successful kelp restoration affects the nutritional landscape and the fitness of kelp forest herbivores. We bridge this knowledge gap with a Before-After-Control-Impact Paired Series (BACIPS) focused on kelp forest restoration where reductions of herbivorous sea urchins in Haida Gwaii resulted in substantial increases in kelp abundance in habitat previously characterized as barrens. Specifically, we document body size specific shifts in the fatty acid (FA) profiles of red sea urchins (*Mesocentrotus franciscanus*) and northern abalone (*Haliotis kamtschatkana*). FAs associated with bacteria and diatoms were elevated in tissues of urchins and abalone in barrens habitat while kelp biomarkers were elevated in restored kelp forest habitat. For urchins, these shifts tracked the increase in gonad mass following kelp forest recovery. For abalone, these results varied depending on animal body size. Specifically, abalone exhibited a continuous size-specific shift from biofilm-associated markers at small sizes to kelp-associated markers as animals increased in size. For both species, a marked increase in essential fatty acids was observed following kelp restoration. Our results demonstrate kelp restoration via sea urchin reduction enhances not only the quantity but also the quality and diversity of food in previously degraded habitats, and subsequently enhances the amount and nutritional quality of roe (i.e., gonads) in sea urchins therein.

## INTRODUCTION

Kelp forests, one of the most productive and diverse ecosystems on earth (Dayton 1985; Steneck et al. 2002), increasingly face compounding threats from climate change and overgrazing. Kelps (i.e., macroalgae in the order Laminariales) span roughly one quarter of the world’s coastal habitats and serve as foundation species for many rocky reef ecosystems (Teagle et al. 2017; Wernberg et al. 2019). By contributing high primary productivity and critical biogenic habitat, kelps form the basis of complex nearshore food webs, increasing secondary productivity and local biodiversity (Smale et al. 2013; Steneck et al. 2013). Additionally, kelp forests provide valuable ecosystem services for human society including the provision of habitat and food for important fisheries, nutrient cycling, and storm protection (Wernberg et al. 2019; Smale et al. 2013). Unfortunately, direct and indirect effects of rapid global change, such as warming trends and extreme heating events (Smale 2020), as well as disruptions to trophic cascades (Estes and Duggins 1995), are reducing the stability and suppressing the resilience of kelp forests worldwide (Wernberg et al. 2019). When sea urchins proliferate, they often graze down macroalgae, resulting in “sea urchin barrens”, and declines in productivity, biodiversity, and fisheries resources (Filbee-Dexter and Scheibling 2014; Graham 2004). Owing to urchins’ extreme resistance to starvation and physiological resilience (e.g., Spindel, Lee, and Okamoto 2021; Dolinar and Edwards 2021), some urchin barrens can persist for decades or more (Filbee-Dexter and Scheibling 2014). Unfortunately, the sea urchins dwelling in such food-depauperate environments tend to have depleted gonads (Harrold and Reed 1985; Kato and Schroeter 1985; Rogers-Bennett et al. 1995; Spindel, Lee, and Okamoto 2021) (i.e., “roe” in a fisheries context), rendering them unviable for either human harvest (Claisse et al. 2013) or as prey (Smith, Tomoleoni, et al. 2021; Eurich et al. 2024). Human interventions to restore kelp forests (sensu Balensiefer et al. 2004) have the potential to ameliorate these harmful impacts (Claisse et al. 2013) and have increasingly shown promising results (Flukes, Johnson, and Ling 2012; Eger et al. 2020; Eger et al. 2022; Layton et al. 2020; Lee et al. 2021; Grime et al. 2023). However, implicit in these results are assumptions of trophic linkages between kelp forest consumers and kelp production (e.g., Graham 2004) that have not been empirically tested (Elliott Smith and Fox 2021).

Understanding how changes in kelp production transfer to consumers is fundamental to kelp forest ecology and conservation, particularly under condition of rapid global change. Food availability is among the strongest drivers limiting individual fitness, population dynamics, and community structure (Okamoto et al. 2012; Mduma, Sinclair, and Hilborn 1999; Olsen et al. 2011; Travis et al. 1987; Levitan 1989; Elliott Smith and Fox 2021; Harrold and Reed 1985) and may determine the degree to which consumers can absorb stressors like warming (Huey and Kingsolver 2019). While the importance of kelp as a biogenic habitat-forming species has been relatively well-documented (Dayton 1985; Graham 2004; Miller et al. 2018), relatively few studies have quantified the use of kelp-derived nutrition by consumers within kelp forests (Elliott Smith and Fox 2021). Importantly, the stable coexistence of competing consumer species in these ecosystems likely depends on some combination of resource partitioning among different species (Schoener 1974) and within a single species as a function of body size (Werner and Gilliam 1984b). Such knowledge of the size specific nature of trophic interactions improves our understanding of the life history of species at risk like abalone and the structure of the community in which it is embedded (Post 2003; Werner and Gilliam 1984a). Diet tracer studies have the potential to resolve some of the uncertainty around the dynamics of such interspecies and/or ontogenetic resource partitioning.

While several approaches have been deployed to trace consumer diets in ecological studies (Nielsen et al. 2018), fatty acid (FA)-based diet tracing provides an attractive compromise between resolution and technical complexity (Jardine, Galloway, and Kainz 2020). This balance is particularly true when assays of wild populations are paired with controlled feeding trials in the laboratory that characterize the modification of dietary FAs during the process of tissue building in focal consumers (Galloway and Budge 2020). Additionally, a FA-based diet tracing approach is well-suited for herbivores like urchins and abalone because macroalgae tend to produce distinct taxa-specific FA signatures (Galloway, Britton Simmons, et al. 2012), and the method has a demonstrated capacity to detect dietary inputs of different macroalgae in laboratory feeding trials (Raymond, Lowe, and Galloway 2014; Schram et al. 2018). Aquaculture studies have demonstrated that specific FAs (e.g., polyunsaturated fatty acids [PUFAs] – fatty acids with two or more double bonds) are essential to the diet of marine consumers because 1) they cannot synthesize certain FAs *de novo*, and 2) they need these compounds for crucial physiological functions (Glencross 2009). Moreover, the availability of PUFAs in the diet can determine the fitness of a wide range of consumers via impacts on reproduction, growth, and survival (Dunning, Russell, and Robker 2014; Fuiman and Perez 2015; Ruiz et al. 2021; Twining et al. 2021). However, the relatively few studies investigating the role of FAs in the performance of marine consumers, particularly *in situ*, stands in stark contrast to the wealth of research focused on other determinants of nutritional quality (e.g., energy content, element ratios, and bulk macronutrients like carbohydrate, protein, and lipid content) (Campanya-Llovet, Snelgrove, and Parrish 2017). Kelp forest restoration presents an opportunity to bridge this knowledge gap. FA-based diet tracing paired with traditional surveys may elucidate how shifting primary production transfers to secondary production and how consumers partition newly available resources in an ecological restoration context.

Here we deploy such a strategy to investigate a Before-After-Control-Impact Paired Series (BACIPS) of kelp forest restoration via reductions of herbivorous sea urchins from 2018 to present in British Columbia, Canada (Lee et al. 2021). The project, called Chiix uu Tll iinasdll (Skidegate Haida language for “Nurturing Seafood to Grow”), involved a collaboration among the Council of the Haida Nation, Parks Canada, Fisheries and Oceans Canada, commercial fishers, academia, and research institutes to restore kelp forests from a barrens state along 3-km of coastline in Gwaii Haanas National Park Reserve, National Marine Conservation Area Reserve, and Haida Heritage Site (hereafter Gwaii Haanas) on X aayda Gwaay (Haida Gwaii). In addition to increasing the abundance and diversity of kelp, our research objectives also included 1) quantifying the degree to which kelp resurgence enhanced the local sea urchin fishery via changes in gonad production, 2) building understanding of how the nutritional seascape changed as a function of depth through the process of kelp recovery, and 3) characterizing how changes in the nutritional seascape were assimilated by co-occurring grazers including endangered northern abalone, *Haliotis kamtschatkana*, and red sea urchin, *Mesocentrotus franciscanus*.

## METHODS

### Kelp forest restoration

The details of the collaborative kelp forest restoration project, Chiix uu Tll iinasdll *Nurturing Seafood to Grow*, co-led by Parks Canada and the Council of the Haida Nation, are published in *Ecological Restoration* (Lee et al. 2021). The initial intensive restoration efforts involving sea urchin removal and *in situ* cracking occurred in fall 2018 and spring 2019 at the restoration site in Gwaii Haanas on X aayda Gwaay, a remote archipelago in northern British Columbia, Canada. The restoration site was located along Gaysiigas Gwaay *Murchison Island* and the control site was along Daa.a Gwaay *Faraday Island*. Sea urchin reduction efforts by Haida Fisheries Program (HFP) and Pacific Urchin Harvesters Association divers spanned from 0-15 m field depth at the restoration site. Haida Fisheries Program and American Academy of Underwater Sciences divers conducted subtidal monitoring surveys on SCUBA along permanent benthic transects to characterize benthic community composition, species abundances, and species size-frequency distributions for invertebrates, macrophytic algae, and fish. Each of the restoration and control sites were divided into two replicate subsites, each of which included two replicate plots. Within each plot, one 30 m long x 2 m wide belt transect was permanently installed along each of three depth isobaths following rocky substrate: 0-2 m (“shallow”), 2-5 m (“mid”), and 5-10 m (“deep”) chart datum depths. Divers surveyed both sites pre-restoration in summer 2017 and 2018, after which almost 90% of sea urchins were manually removed or cracked in-situ in fall 2018 and spring 2019. Post-restoration surveys were conducted annually in the summers of 2019, 2020, and 2021, along with one day of annual restoration maintenance with commercial fishers each spring (Figure 1a & b).

**Figure 1:**
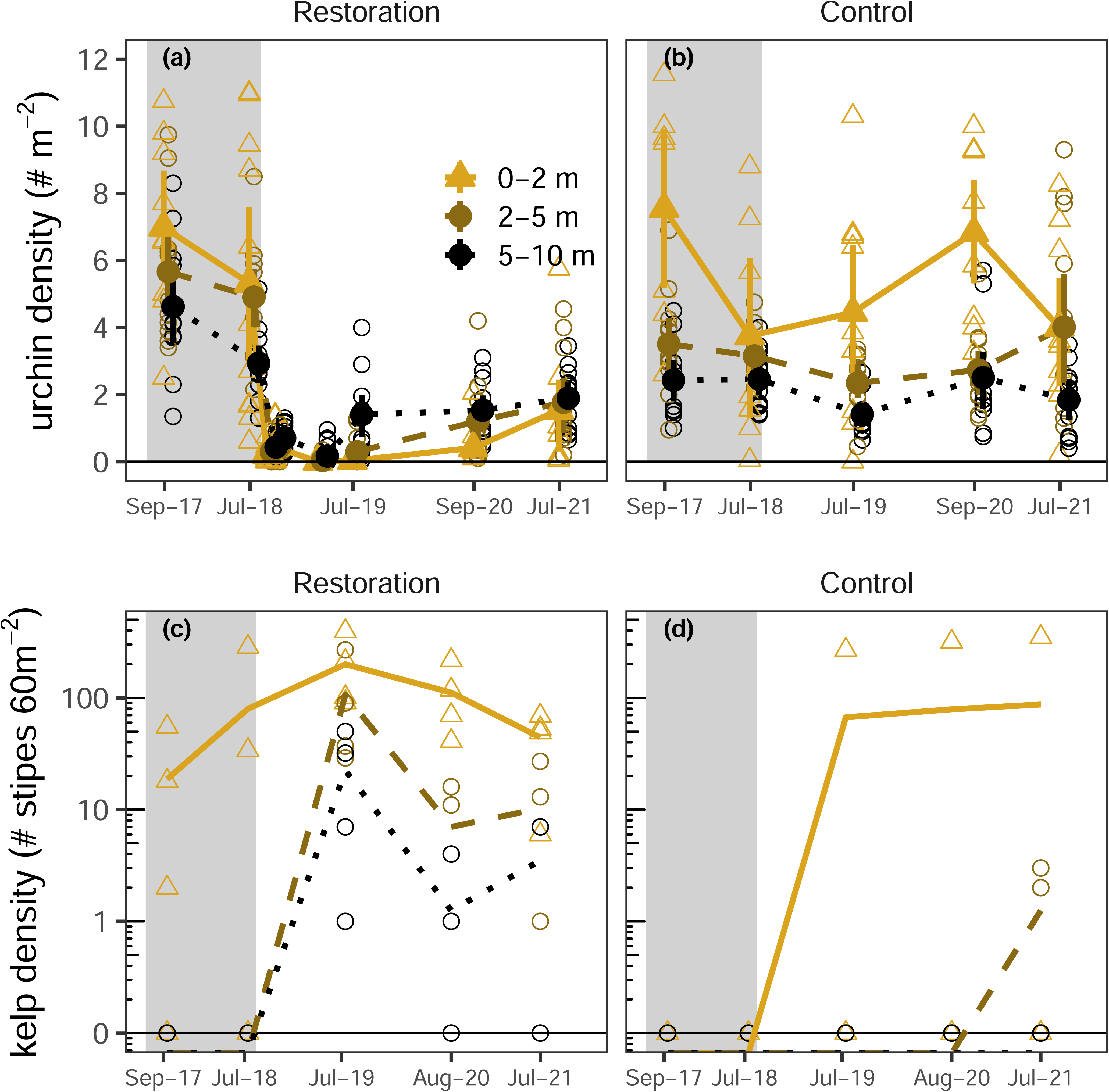
Kelp forest restoration context as part of the Chiix uu Tll iinasdll *Nurturing Seafood to Grow* project (Lee et al. 2021). (**a-d**) Time series illustrating the abundance of red urchin *M. franciscanus* (**a & b**) and kelp (**c & d**) throughout the restoration process as a function of depth zone. (**c & d**) The vertical axis for kelp is on the log scale to spread data more evenly and visualize trends spanning several orders of magnitude. The darker shaded region represents the pre-restoration timeframe, and the lighter region represents post-restoration timeframe. Larger solid symbols and vertical bars represent simple empirical bootstrap means and 95% confidence intervals, respectively, for each depth by survey. Open symbols represent transect level means for each survey.

### Sea urchin gonad assessment

To estimate how shifts in algal food availability translated to shifts in the health and marketability of sea urchins at our study sites, we quantified body size specific gonad mass of the regionally dominant sea urchin species, *M. franciscanus*, in 2018 and 2019 (Figure 1c).

Previous studies have demonstrated that *M. franciscanus* gonad growth is sensitive to food availability (reviewed in Rogers-Bennett and Okamoto 2020). Divers collected benthic stage *M. franciscanus* individuals spanning a representative range of body sizes (28-150 mm test diameter) by hand-picking on SCUBA near, but outside, the shallow and deep permanent monitoring transects, then immediately transferred them to a nearby floating research camp. We measured the test diameter for each individual using precision calipers, cracked the test, measured drained mass, dissected out one of the five gonads for wet mass, and froze samples of a second gonad for fatty acid analysis (see below).

### Abalone condition assessment

We estimated the effects of shifting algal food availability on the health of *H. kamtchatkana* at the same locations and times noted above using a condition index representing the ratio of body mass to volume after accounting for potential biases in sex specific size structure. Divers collected benthic stage *H. kamtschatkana* individuals ranging in body size from 44-125 mm in shell length by hand-picking on SCUBA. We temporarily retained these individuals in flow-through containers to minimize handling stress while collecting non-invasive biometric data and tissue samples. After completing biometrics and tissue sampling, divers carefully returned all individuals to their collection sites by hand on SCUBA. Biometrics included shell length and width, overall body height, wet mass, and sex. We used precision calipers for body dimensions, a digital scale for wet mass, and external visual classification of the gonads for sex. To approximate body volume, V, we applied the geometric equation for an ellipsoid to our metrics for length (*l*), width (*w*), and height (*h*): 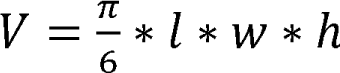. We then modeled wet mass as a function of a log body volume-sex interaction (accounting for potential differences in mass scaling between males and females) plus a treatment-year-depth-sex interaction using a Bayesian regression model with a Gamma likelihood and log link function. We defined the condition index as the intercept of this model for each treatment-year-depth-sex combination.

### Tissue sampling for fatty acid analysis

We used fatty acids as dietary tracers to evaluate how primary consumers assimilate shifts in the abundance and composition of primary production. This approach involved characterizing contemporaneous FA profiles in both primary producers and consumers at the same locations, as well as in a controlled feeding trial in the laboratory (Spindel et al. *in prep*). We biopsied as many algal taxa as possible at our study sites in 2018 (4 classes, 13 orders, 22 families, 39 genera, and 47 species) to generate a herbivore food resource library (sensu Galloway et al. 2015), taking care to manually scrub off conspicuous epibiota from algal thallus tissue prior to storage. We retained voucher specimens and confirmed species identifications for cryptic algal taxa using DNA barcoding (sensu Gabrielson et al. 2023). Our focal consumers included co-occurring, but phylogenetically distant, invertebrate grazers: the gastropod *H. kamtschatkana* and the echinoid *M. franciscanus*. We defined focal sets of biomarkers by reference to well-documented cases in the literature (Galloway, Britton-Simmons, et al. 2012; Kelly and Scheibling 2012) and to our custom algal reference library described above.

To capture signals of potential ontogenetic dietary shifts, we sampled individuals of each species spanning a range of benthic stage body sizes (urchins: 21-133 mm test diameter; abalone: 44-125 mm shell length). We sampled individuals of each species annually in summer 2018 pre-restoration and 2019 and 2020 post-restoration near, but outside, permanent shallow and deep transects at the control and restoration sites (urchins: n = 10-15 per year, site, and depth; after 2018, n = 15 urchins per year, site, and depth; abalone: n = 10-27 per year, site, and depth; after 2018, abalone n ≥ 15 per year, site, and depth). For *M. franciscanus*, we sampled gonad tissue. *H. kamtschatkana* is an endangered species, and as such is protected from lethal sampling, therefore for FA analysis, we collected non-lethal biopsies of epipodia, a practice demonstrated in the laboratory to impart negligible stress for this species (Withler et al. 2001). We used 70% ethanol to sterilize equipment and surfaces between samples to prevent contamination. To prevent degradation of lipids, we immediately froze all harvested tissue samples at -20 in 1.5 mL cryovials then transferred frozen samples to -80 freezers for storage and later lipid extraction and fatty acid composition profiling (see below).

### Fatty acid extraction and analysis

We extracted lipids from tissue biopsies using nonpolar solvents (sensu, Schram et al. 2018; Parrish 1999). First, we lyophilized samples previously frozen at -80 for 48 h to remove all water content and further reduce the probability of lipid degradation prior to analysis. Next, we disrupted and homogenized each tissue sample by grinding it with a pellet pestle. Following homogenization, lyophilized tissue samples were digested in 2 mL chloroform, then sonicated to further disrupt cell membranes, vortexed, and centrifuged (3000 rpm for 5 mins) twice in a 4:2:1 chloroform:methanol:0.9% NaCl solution. After each of these cycles, the lower chloroform layer containing lipids was pipetted out and pooled. We extracted 1 mL of this pooled solution and evaporated it to dryness under N_2_ flow using a nitrogen evaporator (N-Evap, Organomation Associates, Inc). We re-suspended the resulting organic residue in a solution of toluene and 1% H_2_SO_4_:MeOH then sealed samples and transferred them to a heated water bath where they were kept for 90 mins at 90 to transesterify fatty acid methyl esters (FAME). Transesterified FAME solutions were allowed to cool to room temperature before 2% KHCO_3_ was added to neutralize the pH. The resulting FAME mixture was diluted with 2 mL of hexane then vortexed and centrifuged (1500 rpm for 2 mins). The supernatant FAME layer was removed and retained in a pooled sample, and the cycle of hexane addition, vortex mixing, centrifugation, and transfer of the FAME layer to the pooled sample was repeated once more. Following the second cycle, the pooled FAME-hexane solution was evaporated to dryness under N_2_ flow. The resulting FAME residues were re-suspended in 1.5 mL hexane and transferred to 2 mL glass gas chromatograph (GC) vials for analysis on a GC equipped with a DB-23 column (30 × 0.25mm × 0.15µm, Agilent, Santa Clara, CA, USA) using helium as a carrier gas for a GC-mass spectrometer (GC-MS, Model QP2020, Shimadzu Scientific Instruments ©). The heating program employed for the GC-MS analysis followed Taipale et al. 2013. Fatty acids were characterized based on specific ions and relative retention times in the column, then quantified using mass spectra peaks of the major ions following Taipale et al. 2016 using the software package Lab Solutions (Shimadzu Scientific Instruments ©). We then calculated the proportions of different fatty acids based on relative integrated peak areas in the resulting chromatograms.

### Statistical analyses Univariate

We estimated univariate response variables as functions of predictors using Bayesian regression models. To control for uncertainty in model formulae, we conducted model comparison using approximate leave-one-out (loo) cross validation and selected model formulae based on weights estimated using stacking (Vehtari, Gelman, and Gabry 2016; Yao et al. 2018). This method is designed to maximize predictive accuracy of candidate models. In cases where including a body size allometric scaling term increased posterior model weights, we accounted for this effect by including the log of body size (test diameter for urchins and shell length for abalone) as a covariate. Results presented here are from models representing >99% of posterior model weights. We compared parameter posteriors directly within selected models. For animal models, we included a year, treatment (control versus restoration), and depth interaction effect. For the algal model, we included only the effect of taxonomic class. We estimated model posteriors using Stan (Team 2022, 2024b) via the R (Team 2024a) package brms (Bürkner 2017).

We modeled non-compositional and compositional responses using different likelihood distributions and link functions. For non-compositional, continuous, and positive gonad mass responses, we utilized a Gamma distribution and log link function. For compositional fatty acid proportion responses, we implemented a Dirichlet regression. To account for the compositional nature of fatty acid proportions, we modeled a response vector ***Y*** = (*Y*_1_, *Y*_2_,…, *Y_K_*), where each *Y_k_* represents a component fatty acid proportion of the overall profile and the set of component fatty acid proportions sum to one, 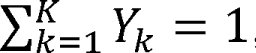, using the Dirichlet distribution, *Y∼Dirichlet(µ, ¢)*. Here, µ = (µ_1_, µ_2_,…, µ_*K*_) is the vector of mean parameters, with each component µ_*k*_ representing the expected value for the corresponding proportion, and ¢ representing a concentration parameter that controls the variability of the distribution. We used a logit link function to constrain predicted proportions within sensible bounds between zero and one and to respect the compositional nature of the data. We used vague priors relative to the predicted scale of responses (Gelman et al. 2013) detailed in the following:

Concentration parameter (¢): Gamma distribution, appropriate given the strictly positive nature of ¢:

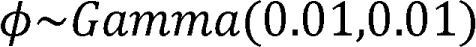

Regression coefficients (/J): Normal distributions associated with each µ_*k*_, with a mean of zero and a standard deviation of 1:

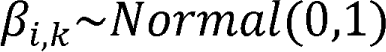

Intercepts: Student’s t distributions were used to offer robustness against outliers:

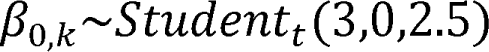

We fit our models using 8,000 iterations across four chains, discarding the first half of the iterations per chain as a warm-up, resulting in a posterior sample of 16,000 iterations for each response, checking that chains converged using visual inspection. For each parameter estimate, we confirmed that R_hat_, the potential scale-reduction factor was less than 1.01 and the minimum effective sample size, n_eff_, was greater than 1,000 (Gelman et al. 2013). To evaluate goodness of fit for our models, we evaluated graphical posterior predictive checks, searching for any systematic differences between model simulations and empirical data (Gelman et al. 2013) and estimated Bayesian *R^2^*values (Gelman et al. 2019).

### Multivariate

We visualized multivariate patterns of fatty acid profiles using ordination and evaluated the significance of predictors using robust distance-based multivariate analysis of variance. We generated non-metric multidimensional scaling ordinations (NMDS) (Clarke 1993) based on Aitchison dissimilarity, which utilizes a centered log ratio transformation to account for compositionality (Aitchison 1982). NMDS ordinations were fit using the package vegan (Oksanen et al. 2007) in R (Team 2024a). Ordination stress is reported within each NMDS plot. We fit correlation vectors for constituent fatty acids, such that vector lengths were scaled by correlation coefficients with respect to NMDS ordination scores for each data point. Fitted fatty acid vector directions indicated the ordination space toward which the focal fatty acid changed most rapidly and had the greatest correlation with the ordination configuration (Oksanen et al. 2023). We ranked the contributions of individual fatty acids to multivariate dissimilarity between groups of interest using a SIMPER analysis (Clarke 1993) using the package vegan (Oksanen et al. 2007) in R (Team 2024a). To assess significance of predictor variables on multivariate responses while accounting for heteroskedasticity and unequal sample sizes, we ran W* tests (Hamidi et al. 2019). We ran post hoc pairwise comparisons using distance-based Welch *t*-tests, T^2^ (Alekseyenko 2016), correcting for multiple comparisons using a Bonferroni adjustment.

## RESULTS

### Restoration context

Manual removal and *in situ* cracking of sea urchins in fall 2018 and spring 2019 substantially reduced local densities of *M. franciscanus* at treatment sites (Figure 1a) and the following summer yielded many-fold increases in subtidal kelp densities (Figure 1c) along with increased gonad mass among remaining *M. franciscanus* individuals inhabiting deep transects (Figure 1g) (Lee et al. 2021). Specifically, these sea urchin reduction efforts reduced densities of *M. franciscanus* by approximately 88%, going from 49.36 individuals/10 m^2^ (± 4.38 SE) in summer 2018 pre-restoration surveys to 5.76 individuals/10 m^2^ (± 1.22 SE) post-restoration in summer 2019 (Figure 1a). During the same timeframe, *M. franciscanus* densities remained relatively consistent at the control site (Figure 1b). Prior to the urchin reductions, deep transects were urchin barrens, habitats denuded of macrophytic algae by overgrazing sea urchins. The sea urchins dwelling in these barrens were emaciated and possessed diminutive gonads (sensu Spindel, Lee, and Okamoto 2021). In the post-restoration 2019 summer survey, bull kelp, *Nereocystis luetkeana*, increased by approximately 6687%, going from 0.83 stipes/60 m^2^ (± 0.78 SE) to 56.33 stipes/60 m^2^ (± 32.26 SE) (Lee et al. 2021) at the restoration site (Figure 1c). One shallow transect at the control site coincidentally had comparable kelp density to the restoration site from July 2019 onward, suggesting generally higher recruitment that summer (Figure 1d). However, all other transects at the control site during the same timeframe had at most 2-3 stipes/60 m^2^ and mostly no kelp at all (Figure 1d).

### Consumer body condition

Body condition improved in both grazer taxa at the restoration site from 2018 pre-restoration to 2019 post-restoration (Figure 2). After correcting for body size, gonad mass among remaining deep *M. franciscanus* individuals at the restoration site increased approximately 89% from 2018 pre-restoration to 2019 post-restoration (Figure 2c; 2018 μ = 7.517 g, 95% highest probability density interval [HPD] = 6.493 - 8.738 g; 2019 μ = 14.210 g, 95% HPD = 12.253 – 16.620 g) in alignment with the resurgence of deep kelp at the restoration site after the urchin reductions. Gonad mass generally increased exponentially with body size (i.e., test diameter; Figure 2a-d). Over the same timeframe, *H. kamtschatkana* body condition index (BCI) generally increased at the restoration site, particularly for females (Figure 2e, g). On average, female BCI at the restoration site increased by 28.4% (95% HPD = 14.8-45.3%) in shallow individuals and 30.9% (95% HPD = 7.5-53.8%) in deep individuals. In contrast, female BCI at the control site increased only marginally in shallow individuals and marginally decreased in deep individuals (Figure 2f, h; shallow: 13.4%, 95% HPD = -9.5-31.5%; deep: -1.7%, 95% HPD = -18.6-16%).

**Figure 2:**
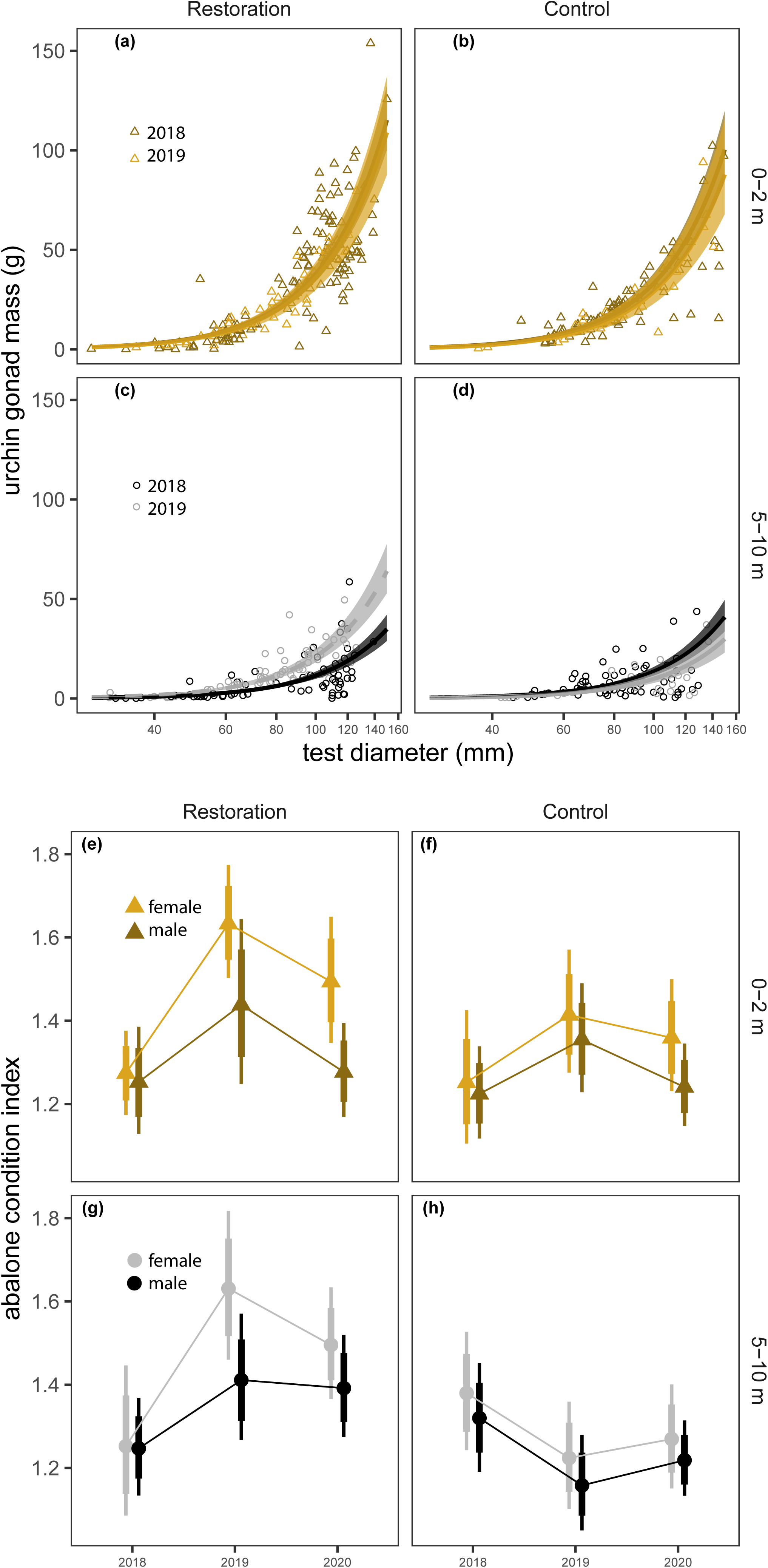
Body condition assays for (**a-d**) *M. franciscanus* and (**e-h**) *H. kamtschatkana.* (**a-d**) Gonad mass as a function of body size, depth, and treatment, pre- and post-restoration. The horizontal axis is on the log scale to highlight that modeling the relationship between gonad mass and body size was more accurately modeled allometrically as a power function, as opposed to isometrically as a linear function. Lines and shaded ribbons represent conditional modeled means and 95% highest probability density intervals, respectively. Open symbols represent empirical data points. Dashed lines represent instances with significant year over year differences in body size specific gonad mass for a given depth. (**e-h**) Condition index representing the ratio of body wet mass to volume as a function of sex, depth, and treatment, pre- and post-restoration after accounting for body size (i.e., shell length). Points represent conditional modeled means and thin and thick vertical bars represent 95%, and 80% highest probability density intervals, respectively.

### Fatty acid biomarkers

#### Echinoid grazer: Mesocentrotus franciscanus

In general, the gonad FA profile of deep-dwelling *M. franciscanus* individuals at the restoration site transformed from a barrens type to a kelp forest type after restoration efforts (Figure 3a-f). There was a significant three-way interaction effect of year, treatment, and depth on multivariate FA profiles (W* = 20.388, p = 0.001, nrep = 999). Neither the restoration nor the control sites differed year-over-year in shallow habitats (restoration 2019-2020: T^2^ = 0.942, W* p = 0.576, nrep = 999; control 2019-2020: T^2^ = 0.781, W* p = 0.767, nrep = 999). However, profiles differed by treatment and depth (W* = 17.778, p = 0.001, nrep = 999). Profiles of deep versus shallow individuals at the restoration site differed in 2018 pre-restoration (T^2^ = 16.742, W* p = 0.001, n_deep_ = 13, n_shallow_ = 12), but not post-restoration (2020: T^2^ = 1.918, W* p = 0.264, n_deep_ = 15, n_shallow_ = 15). In contrast, individuals at the control site differed by depth across all years (W* p ≤ 0.001).

**Figure 3:**
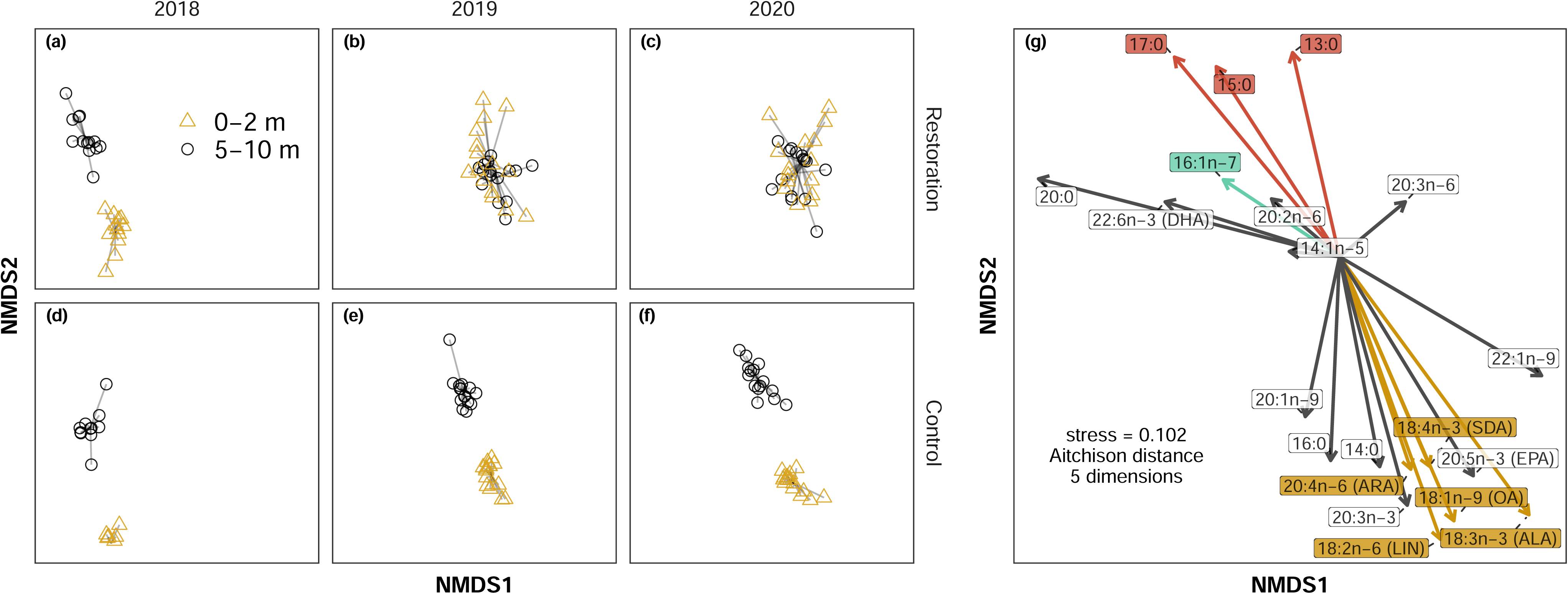
Fatty acid profile change in sea urchins in response to kelp restoration and non-metric multidimensional scaling (NMDS) ordination plots illustrating dissimilarity between multivariate fatty acid profiles. (**a-f**) Time series of multivariate fatty acid profile shifts as a function of depth and treatment from pre-restoration (2018) to post-restoration (2019 & 2020). Points represent measured profiles comprised of 34 fatty acids. (**g**) Arrows indicate fitted correlation vectors for the top twenty most influential constituent FAs driving multivariate dissimilarity, where each arrow’s length is scaled as a function of its correlation with the ordination scores for each data point. Specific vectors are highlighted in color to emphasize the contribution of key classes of biomarkers to dissimilarity: kelp (gold), bacteria (red), and diatoms (aquamarine).

Much of the dissimilarity in FA profiles between shallow and deep animals was driven by two classes of FAs, specifically those compounds typically associated with kelp and those associated with bacteria and diatoms (i.e., biofilm; Figure 3g). The summed proportions of kelp biomarker FAs in *M. franciscanus* gonads (oleic acid [OA], alpha-linolenic acid [ALA], alpha-linoleic acid [LIN], stearidonic acid [SDA], arachidonic acid [ARA], and vaccenic acid) increased along deep transects post-restoration at the restoration site (Figure 4a), in alignment with the resurgence of deep kelp abundance and concomitant increase in gonad mass per individual urchin at the restoration site over the same period (Figure 2c). The summed proportion of kelp biomarkers in gonad tissue increased by approximately 27% along deep transects in the first year post-restoration at the restoration site, and subsequently maintained a similar proportion the following year (Figure 4a; 2018 μ = 0.200, 95% highest probability density interval [HPD] = 0.178 - 0.221; 2019 μ = 0.255, 95% HPD = 0.232 - 0.279; 2020 μ = 0.291, 95% HPD = 0.269 – 0.314). In contrast, the proportions of these FAs did not change significantly at the control site over the same period (Figure 4b; 2018 deep μ = 0.180, 95% HPD = 0.159 - 0.201; 2019 deep μ = 0.160, 95% HPD = 0.140 - 0.180). The proportion of nutritionally valuable PUFAs in deep urchins at the restoration site increased to levels comparable with shallow urchins post-restoration (Figure 4c), whereas this proportion was consistently lower in deep individuals relative to shallow individuals at the control site (Figure 4d).

**Figure 4:**
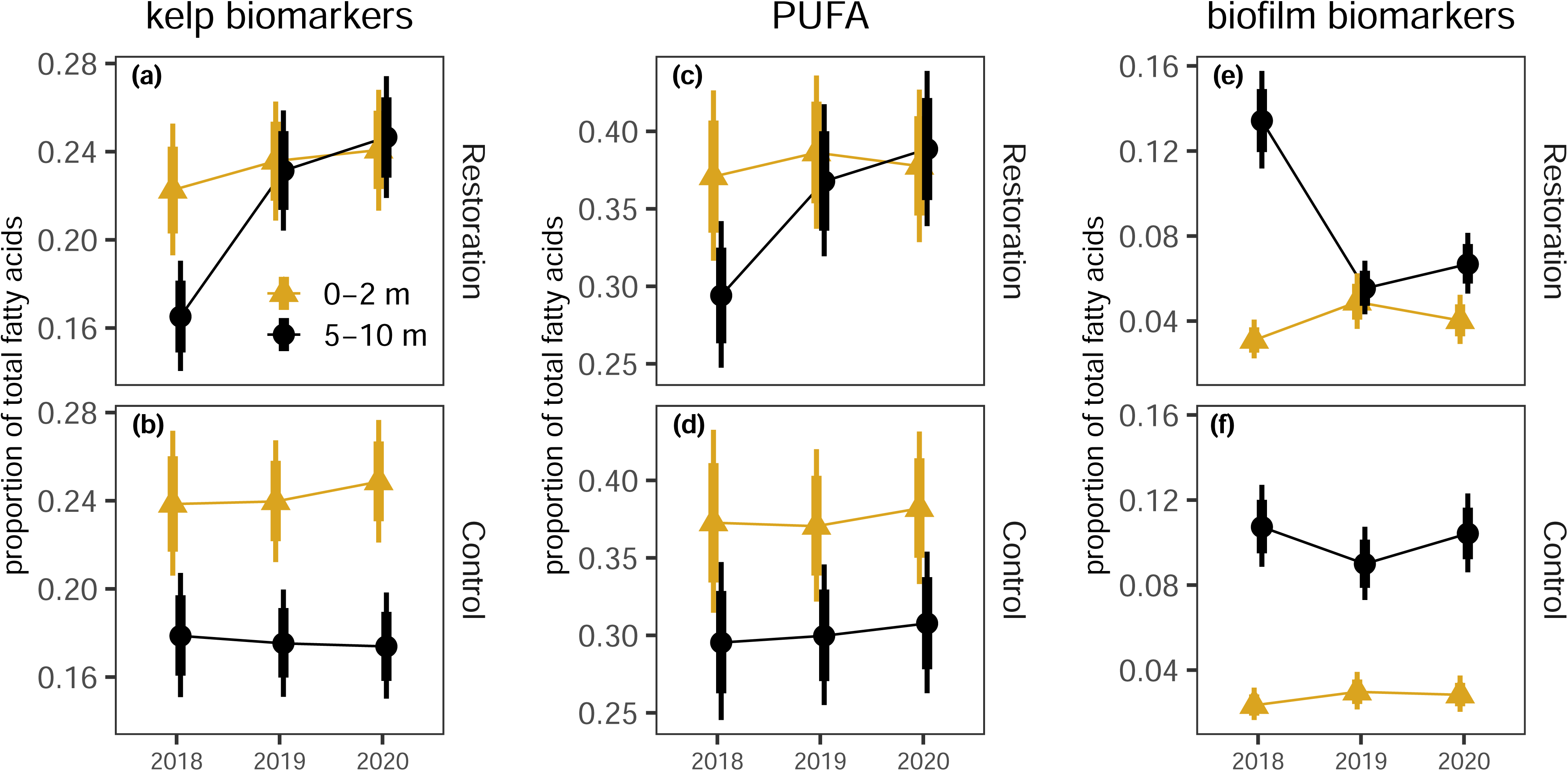
Univariate comparisons of well-documented fatty acid biomarkers in red sea urchin gonad tissue as a function of depth and treatment. (**a & b**) Kelp biomarker, (**c & d**) kelp biomarker plus nutritionally valuable poly-unsaturated fatty acids (PUFA - fatty acids with two or more double bonds), and (**e & f**) biofilm biomarker proportions in sea urchins following kelp restoration. Points represent conditional modeled means for the summed proportions by biomarker category and thin and thick vertical bars represent 95%, and 80% highest probability density intervals, respectively.

The other class of FAs that consistently differentiated gonad FA profiles by depth was biofilm biomarkers (Figure 4e, f). At the control site, individuals from deep, barrens habitat had higher proportions of FAs with putative origins in bacteria and diatoms (Dalsgaard et al. 2003; Kelly and Scheibling 2012). Specifically, the proportions of odd-chain FAs 13:0, 15:0 and 17:0, generally considered to be bacterial metabolites, and 16:1n-7, palmitoleic acid, generally associated with diatoms, were consistently higher in deep versus shallow habitats at the control site (Figure 4f). In contrast, deep-dwelling urchins at the restoration site initially had higher proportions of these FAs than their shallow counterparts in 2018, but these proportions decreased to a level comparable to shallow urchins in 2019 post restoration and remained at similar proportions the following year (Figure 4e).

#### Gastropod grazer: Haliotis kamtschatkana

Kelp biomarkers consistently increased in *H. kamtschatkana* in deep habitats at the restoration site post-restoration (Figure 5, 6), but other FA profile shifts were more nuanced than in *M. franciscanus*, owing in part to a strong effect of body size (Figure 6c, d). Specifically, the proportion of biofilm biomarkers in *H. kamtschatkana* epipodia decreased as a function of body size with slightly higher proportions in small deep-dwelling individuals (Figure 6c). Conversely, the proportion of kelp biomarkers increased as a function of body size with consistently higher proportions in shallow-dwelling individuals for all but the largest body sizes recorded (Figure 6d). Like *M. franciscanus* gonads, multivariate FA profiles in *H. kamtschatkana* epipodia also had a significant three-way interaction effect of year, treatment, and depth (Figure 5a-f; interaction effect W* = 29.527, p = 0.001, nrep = 999). The proportion of kelp biomarkers increased in epipodia tissue by approximately 27% along deep transects the first year post-restoration at the restoration site, then again by an additional 14% the following year (Figure 5a; restoration site: 2018 μ = 0.200, 95% HPD = 0.178 – 0.222; 2019 μ = 0.255, 95% HPD = 0.232 – 0.279; 2020 μ = 0.291, 95% HPD = 0.269 – 0.314). Additionally, the proportion of kelp biomarkers in abalone along deep transects at the restoration site were approximately 59% higher than in abalone at the control site at similar depth in the first year post-restoration, and approximately 47% higher the following year. (Figure 6a, b; control site: 2018 μ = 0.180, 95% HDP = 0.159 - 0.201; 2019 μ = 0.160, 95% HDP = 0.140 - 0.180; 2020 μ = 0.198, 95% HDP = 0.180 - 0.216).

**Figure 5:**
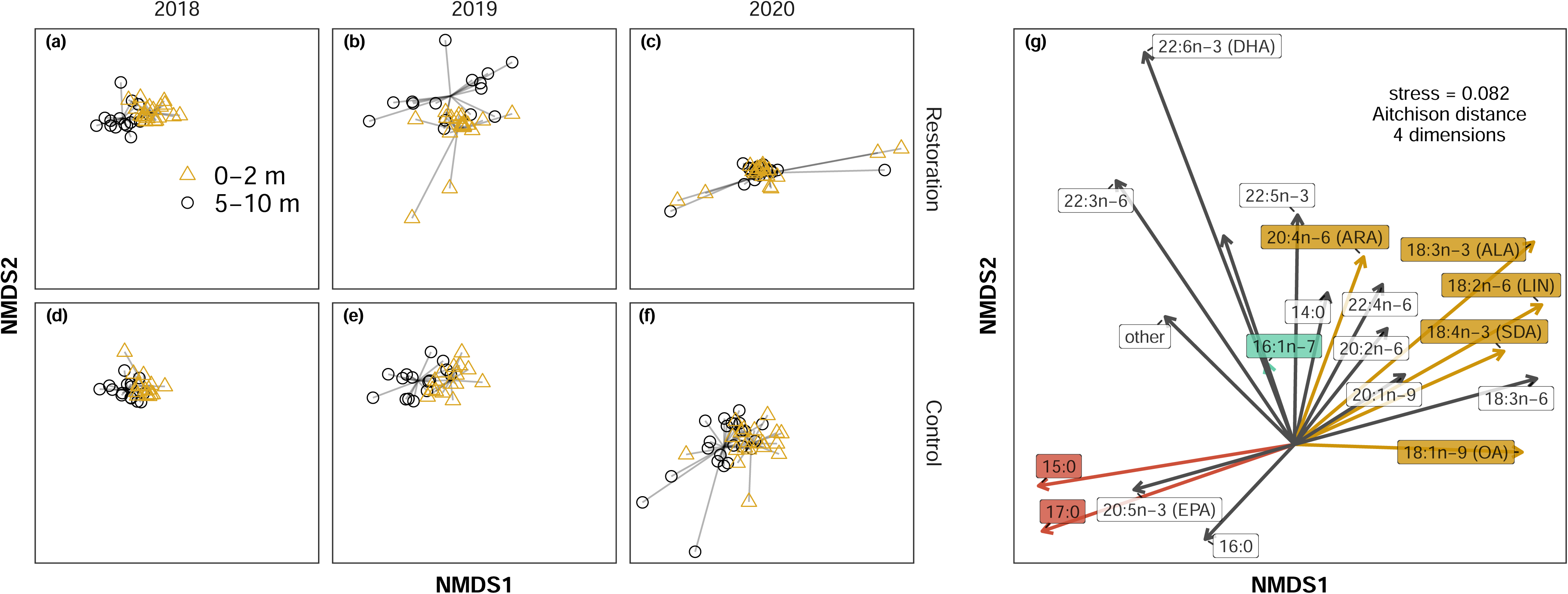
Fatty acid profile shift in abalone in response to restoration and non-metric multidimensional scaling (NMDS) ordination plots illustrating dissimilarity between multivariate fatty acid profiles. (**a-f**) Time series of multivariate FA profile shifts as a function of depth and treatment from pre-restoration (2018) to post-restoration (2019 & 2020). Points represent measured data points comprised of 34 fatty acids. (**g**) Arrows indicate fitted correlation vectors for the top twenty most influential constituent FAs driving multivariate dissimilarity, where each arrow’s length is scaled as a function of its correlation with the ordination scores for each data point. Specific vectors are highlighted in color to emphasize the contribution of key classes of biomarkers to dissimilarity: kelp (gold), bacteria(red), and diatoms (aquamarine).

**Figure 6:**
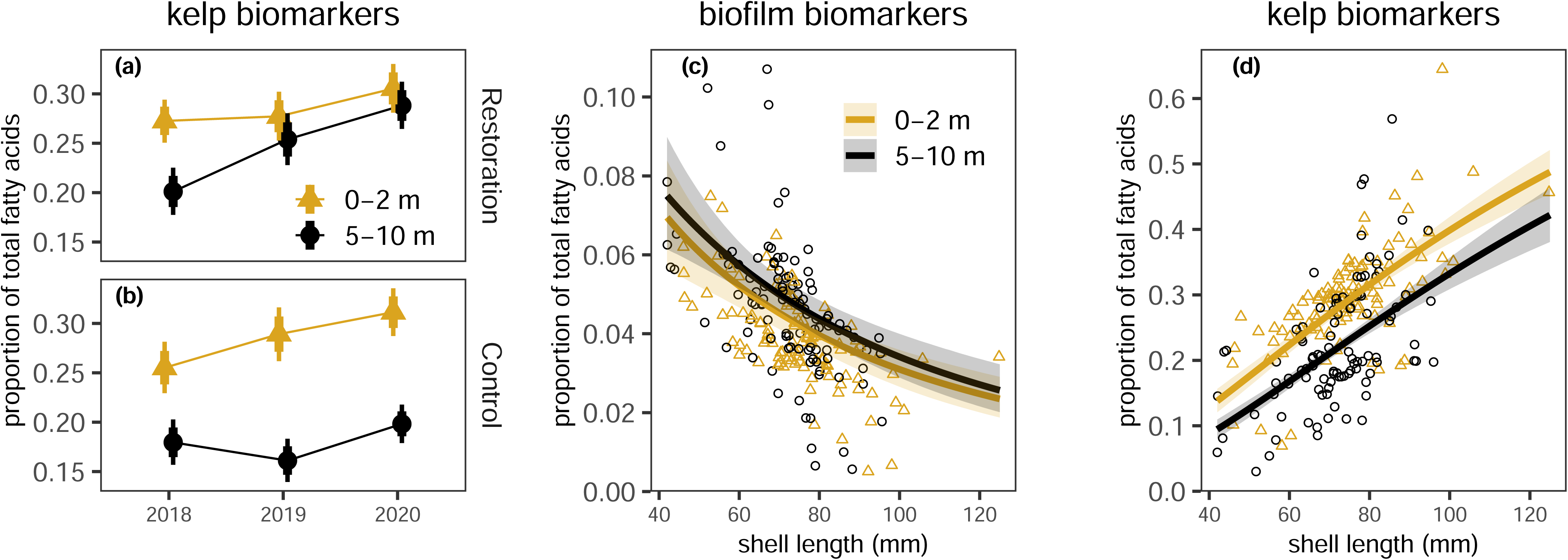
(**a & b**) Comparison of kelp biomarkers in abalone as a function of depth and treatment. Points and thin and thick vertical bars represent conditional modeled means, 95%, and 80% highest probability density intervals [HPD], respectively, pre-restoration (2018) and post-restoration (2019 & 2020). (**c & d**) Relationship between body size and assimilation of (**c**) biofilm versus (**d**) kelp biomarkers. Lines and faded ribbons represent conditional modeled means and 95% HPD, respectively. Faded points represent empirical data points. Horizontal axis is on the log scale to spread data more evenly and improve the visibility of overall trends.

## DISCUSSION

Overgrazing threatens kelp forests worldwide, particularly in the context of rapid global change. While restoration efforts involving sea urchin reductions have often resulted in local gains in kelp abundance, explaining how such gains transfer to resident consumers has involved untested assumptions regarding trophic linkages. In this study, we show that when kelp abundance increases in response to restoration (Figure 1), primary consumers assimilate distinctive biomarkers of kelp, most notably in deeper habitats previously devoid of macroalgae in alignment with improvements in body condition (Figure 2). This trophic transfer yields sea urchin roe with nutritional quality comparable to that found in rich shallower waters.

Additionally, biofilm biomarkers tended to distinguish consumers dwelling in barrens habitat. However, assimilation of kelp and biofilm FAs was dependent on body size in the more mobile consumer species, *H. kamtschatkana* (Figure 6). These results lead us to three key general conclusions. First, the same traits that give *M. franciscanus* the capacity to suppress the recovery of kelp forests from barrens states (i.e., starvation resistance and resilience) also make them good candidates for conservation aquaculture (sensu Angwin, Hentschel, and Anderson 2022; Froehlich, Gentry, and Halpern 2017) and presents targeted urchin reductions as a potentially effective tool for roe enhancement for fisheries markets. This insight stems from the results of our kelp forest restoration project, Chiix uu Tll iinasdll (Lee et al. 2021), in context with our laboratory feeding experiment (Spindel et al *in prep*). The long-term success of applying this tactic would potentially require tuning the magnitude of urchin reductions to simultaneously achieve both adequate kelp recovery and a sustainable urchin fishery. Quantifying effective target densities that achieve these parallel goals and building understanding of how different environmental factors modulate these targets are important future research objectives. Second, FA biomarkers suggest *H. kamtschatkana* undergoes an ontogenetic dietary shift from biofilms to kelp as they grow larger (Figure 6). Third, kelp forest restoration by sea urchin reduction can subsidize the nutritional seascape of deeper habitats and extend the depth of kelp forest habitats, also generating refugia from predation, fishing, and stressful abiotic conditions for benthic species at risk such as *H. kamtschatkana*.

Sea urchins can buffer the detrimental effects of fluctuations in food availability by expressing highly plastic metabolic states. For example, the tropical *Diadema antillarum* physically shrinks under food deprivation, reducing its metabolic demands and increasing survivorship and reproductive output, and conversely grows rapidly when food is abundant (Levitan 1989). This life history strategy has the apparent effect of stabilizing populations near carrying capacity (Levitan 1989). Though plasticity in body size (i.e., test diameter) in temperate taxa like *M. franciscanus* is more dubious (Ebert 2004; Smith and Garcia 2021), other forms of metabolic plasticity like variable allocation of energy towards storage in gonads (Smith and Garcia 2021; Dolinar and Edwards 2021; Okamoto 2014; Rogers-Bennett and Okamoto 2020) and flexible resting metabolic rate (Spindel, Lee, and Okamoto 2021) seem to offset starvation mortality and facilitate resilient reproductive output. Similarly, we show that, after accounting for body size, the gonad mass of *M. franciscanus* increased by 89% in deeper habitat in alignment with large gains in deeper kelp abundance one year after targeted sea urchin reductions. Similar responses to sea urchin reductions have been shown in California with the recovery of kelp in overgrazed barrens (Grime et al. 2023). Together with evidence from laboratory experiments (Angwin, Hentschel, and Anderson 2022; Okamoto 2014; Rogers-Bennett and Okamoto 2020, Spindel et al *in prep*), these results suggest *M. franciscanus* is also a suitable candidate for conservation aquaculture in regions where they are hyperabundant in barrens, adding to a portfolio of strategies for ecosystem-based management of marine species. Furthermore, our FA-based diet tracing approach showed that in addition to an increase in gonad mass among remaining *M. franciscanus*, targeted urchin reductions and subsequent kelp recovery also augmented the nutritional quality of their roe, further enhancing their value to urchin fisheries as food for people.

Fatty acids also elucidated how restoration-driven shifts in food availability affect the trophic ecology of more mobile gastropod grazers like *H. kamtschatkana*, which compete for overlapping food sources with *M. franciscanus*. Because the proportions of biofilm FAs decreased while the proportions of kelp FAs increased as a function of body size, we can infer that this species undergoes an ontogenetic dietary shift. Previous research on a congener, *H. discus hannai*, inferred a similar shift using stable isotope analysis (Won et al. 2010). Such insight helps explain reports of ontogenetic shifts in habitat association with respect to depth for *H. iris* (e.g., Aguirre and McNaught 2012). *H. kamtschatkana* inhabiting sea urchin barrens often have restricted access to kelp, limited to shallow rocky refugia that inhibit sea urchin herbivory because of exposure to higher disturbance from wave energy (Lee et al. 2016). However, these refugia from sea urchins come at the cost of greater risk of mortality from predators, human harvest, and environmental variability. It is far easier for intertidal and diving animal predators as well as human fishers to access abalone in the shallows than deeper in the subtidal. Moreover, the shallows are more susceptible to extreme changes in abiotic conditions (e.g., marine heatwaves, [MHW]), increasing the frequency and magnitude of physiological stress. Unlike sea urchins, which tend to be more resistant to warming (Byrne et al. 2011, Karelitz et al. 2017), acidification (Uthicke et al. 2016), starvation (Spindel, Lee, and Okamoto 2021), and other stressors, abalone are potentially more vulnerable due to lower metabolic flexibility and body size specific dietary needs. For example, recent climate-driven collapses in kelp forests in northern California, USA, drove red abalone out of safer cryptic habitats toward riskier shallows in search of food (Rogers-Bennett and Catton 2022). This shift in habitat use not only exposed red abalone to more stressful environmental conditions, but also likely resulted in a misleading perception of plenty among fishers, leading to overharvest and ultimately the collapse of the fishery in 2018 (Rogers-Bennett and Catton 2022). Our dietary tracers as well as previous studies on a congener (Won et al. 2010) suggest that whereas juvenile abalone may have the option to subsist on biofilms in safer, deeper, more cryptic habitats, larger breeding adults likely require macrophytic algae in the class Phaeophyceae which includes kelp (Figure 6). In a context of increasingly frequent and intense climatic extreme events, traditional proactive management strategies like marine protected areas, fishing gear restrictions, and catch size limits, may therefore be inadequate to effectively recover abalone populations. Our results show that investing in targeted sea urchin reductions designed to recover deeper kelp can improve body condition in resident abalone, particularly for crucially important female breeders and may extend access to kelp for abalone into deeper, less metabolically demanding and higher-risk habitat. By extension, we would predict that other kelp grazer taxa may gain similar depth refugia, but this prediction should be tested with further empirical research expanding analyses to other feeding guilds and higher trophic levels.

This BACI study and its accompanying laboratory experiment (Spindel et al *in prep*) help explain the trophic transfers of kelp forest recovery to grazers. Additionally, our results generate plausible, testable hypotheses for how benthic communities are reorganized as a result of the dynamics of these trophic transfers. How such effects might be modulated by global change drivers like ocean warming is an important future research objective, particularly because food availability may constrain the degree to which marine ectotherms can tolerate the added metabolic demands of gradual warming, acidification, low oxygen, and extreme heating events (i.e., marine heatwaves). Moreover, the degree to which primary consumers depend on kelp versus other producers such as phytoplankton remains a point of contention (Duggins and Eckman 1997; Miller and Page 2012; Docmac et al. 2017), which is consequential for ecosystem-based management strategy. Resolving this question may require the application of higher resolution diet tracers such as compound-specific stable isotopes (Elliott Smith and Fox 2021). In our study, other FAs also contributed to consistent depth-specific differences, but their putative sources are more ambiguous (e.g., saturated FAs [SAFAs] – fatty acids with no double bonds) therefore we opted to constrain our conclusions to a more conservative subset for which we have relatively high confidence in their sources. Examining ratios of specific FAs (e.g., 16:1n-7/20:5n-3, an indicator of diatoms (Kelly and Scheibling 2012)) could help resolve fine-scale diets within the broader category of biofilm biomarkers reported here, which may be particularly important for dietary specialists as opposed to generalists. Our results focus on two key consumer taxa among many in a dynamic community. The degree to which other consumer taxa respond similarly to kelp restoration is an important subject for future research.

Overgrazing accounts for a large proportion of kelp deforestation worldwide (Steneck et al. 2002), and such habitat degradation has cascading negative impacts on nearshore ecosystems and the critical services they provide (Graham 2004; Rogers-Bennett and Catton 2019). This phenomenon has been exacerbated by recent climatic trends (Smale 2020; Smith, Burrows, et al. 2021) and the extirpations of key predators (Burt et al. 2018; Estes and Palmisano 1974; Watson and Estes 2011) that otherwise regulate sea urchin populations. Targeted sea urchin reduction is emerging as an effective technique for increasing the abundance of kelp (Lee et al. 2021; Miller, Blain, and Shears 2022) and enhancing the fishery value of resident taxa (Grime et al. 2023).

Such intervention may best be applied as part of a multifaceted ecological restoration strategy involving, in addition, promoting the recovery of natural predators, marine protected areas, and conservation aquaculture (Eger et al. 2022; Miller, Blain, and Shears 2022). This study is among the first to document how recovering kelp transfers valuable nutrients to resident consumers.

Moreover, this study demonstrates the value of pairing observational ecological surveys with diet tracing using molecular techniques when, as is often the case (Nakazawa 2015), consumers shift their diet throughout ontogeny and organize their habitat associations and interactions with conspecifics and other taxa accordingly.

## ACKNOWLEDGEMENTS

Haawa *thanks* to the Gwaii Haanas Archipelago Management Board and Council of the Haida Nation for their support of this restoration initiative. We thank Michael Thomas for assistance with gas chromatography-mass spectrometry for fatty acid analysis at the Oregon Institute of Marine Biology. Much appreciation for in-kind SCUBA diving capacity from Fisheries and Oceans Canada and Hakai Institute for subtidal surveys. This study was funded by an FSU APACT grant to DKO, Parks Canada Conservation and Restoration (CoRe) project (no. 1808) funding to Gwaii Haanas National Park Reserve, National Marine Conservation Area Reserve, and Haida Heritage Site with LL as technical lead and DKO via contribution agreement with FSU, NSF grant 2023649 to DKO, the William R. and Lenore Mote Eminent Scholar in Marine Biology Endowment at FSU to NBS, a Professional Association of Diving Instructors (PADI) Foundation Research Grant to NBS, an Academy of Underwater Arts and Sciences Zale Parry Scholarship to NBS, and a Phycological Society of America (PSA) Grant-in-Aid of Research to NBS.

## CONFLICT OF INTEREST STATEMENT

The authors declare no conflicts of interest.

## Open Research statement

Data and/or code are provided as private-for-peer review on the Zenodo digital archive (DOI:*pending acceptance or invitation to revise*).

## Notes

### Competing Interest Statement

The authors have declared no competing interest.

